# Benchtop Carbon Fiber Microelectrode Array Fabrication Toolkit

**DOI:** 10.1101/2021.03.22.436422

**Authors:** Julianna M. Richie, Paras R. Patel, Elissa J. Welle, Tianshu Dong, Lei Chen, Albert J. Shih, Cynthia A. Chestek

## Abstract

**Background:** Conventional neural probes are primarily fabricated in a cleanroom, requiring the use of multiple expensive and highly specialized tools.

**New method:** We propose a cleanroom “light” fabrication process of carbon fiber neural electrode arrays that can be learned quickly by an inexperienced cleanroom user. This carbon fiber electrode array fabrication process requires just one cleanroom tool, a parylene-c deposition machine, that can be learned quickly or outsourced to a commercial processing facility at marginal cost. Our fabrication process also includes hand-populating printed circuit boards, insulation, and tip optimization.

**Results:** The three different tip optimizations explored here (Nd:YAG laser, blowtorch, and UV laser) result in a range of tip geometries and 1kHz impedances, with blowtorched fibers resulting in the lowest impedance. While previous experiments have proven laser and blowtorch electrode efficacy, this paper also shows UV laser cut fibers can record neural signals *in vivo*.

**Comparison with existing methods:** Existing carbon fiber arrays either do not have individuated electrodes in favor of bundles or require cleanroom fabricated guides for population and insulation. The proposed arrays use only tools that can be used at a benchtop for fiber population.

**Conclusions:** This carbon fiber electrode array fabrication process allows for quick customization of bulk array fabrication at a reduced price compared to commercially available probes.

## Introduction

Much of neuroscience research relies upon recording neural signals using electrophysiology (ePhys). These neural signals are crucial to understanding functions of neural networks and novel medical treatments such as brain machine interfaces [1]–[3]. Research is conducted using easy-to-assemble probes or commercially available neural recording electrodes. Neural recording electrodes, unique tools with micron-scale dimensions and fragile materials, require a specialized set of skills and equipment to fabricate. A variety of specialized probes have been developed for specific end uses (e.g. a target region) and experiments. However, experiments must either be designed around currently available commercial probes or a lab must invest in the development of a specialized probe which is a lengthy process. Due to the wide variety of neural research, there is high demand for a versatile ePhys probe [1], [4]–[6]. An ideal ePhys probe would feature a small recording site, low impedance [7], and customizable geometry to adapt to the experimental target regions.

Traditionally, commercial probes are silicon-based due to their ability to be fabricated in large batches with a high recording site density [8], [9] and relatively reliable functional stability [10]–[13]. Silicon arrays typically come in one of two designs: a “bed of needles” (Utah) or a planar shank (NeuroNexus [14], Cambridge Neurotech [15], Neuropixel [16]). If an experiment requires a more specific probe geometry than what is commercially available it would require an intensive design, fabrication, and validation process [4], [17], [18]. However, both commercial designs cause tissue damage [19], [20], continuous inflammation around shank sites, increased immune cell response, and decreased neural body density [20]–[22]. The silicon based electrodes corrode and crack [12], [23], insulation detaches or dissolves[10], [24], and traces break from internal and external stresses [12], [23], [25]. These issues cause signals to be lost either due to the mechanical separation of the electrode from the backend connector or in the resulting noise increase that accompanies these failures. Additionally, commercial probes are expensive to produce. The high cost of these probes limits the number of experiments that can be done at a given time. With cost and biocompatibility issues as deterrents, labs often turn to fabricating their own probes with alternative methods and materials.

Thus, sub-cellular scale neural probes are being developed to improve the biological response. Net10/50 probes [5], [26] and silicon carbide “ultramicroelectrodes” [24] use thin flexible shanks to reduce insertion damage. The probes insert quickly and show reliable recordings. The probes also tend to be more mechanically robust due to the flexible nature of the cable reducing the stresses on the electrode sites compared to a traditional silicon array. It is also easy to create a batch of this type of probes at once. However, these devices require either an additionally fabricated and integrated shuttle [5], [26] or need to be coated with a stiffening agent for insertion [24]. Both options offer low damage insertion with neural spike recording capability, but are not commercially available. Fabrication of either design is very difficult to replicate without specialized cleanroom facilities [5], [27], [28] and experience with fabrication processes.

To avoid the expense and difficulty of manufacturing state-of-the-art devices, many labs make their own microwire arrays. Microwires are customizable at the benchtop with reliable results or can be purchased commercially (e.g. Microprobes for Life Science, Tucker-Davis Technologies), offering low damage insertion, a wide recording area, and a small electrode site. However, microwires deform upon insertion [29], cause tissue damage, and can deviate from their initial target [19], [29]–[31]. Commercial microwires have caused moderate inflammation and encapsulation, degrading the electrode [6], [32], [33]. The microwires may also corrode *in vivo*, causing cracks in the insulation and electrode sites [30]. In addition, microwire arrays often require a headstage that limits which small animal models can be studied to those at least the size of a mouse. Commercial headstages can require additional cleanroom processing to improve back end connectorization [34].

In response to the high cost and biocompatibility issues, carbon fiber electrodes may offer an avenue for neuroscience labs to build their own probes without the need for specialized equipment. Carbon fibers are an alternative recording material with a small form factor that allows for low damage insertion. Carbon fibers provide better biocompatibility and considerably lower scar response than silicon [35]–[37] without the intensive cleanroom processing [29]. Carbon fibers are flexible, durable, easily integrated with other biomaterials [37], and maintain electrode integrity *in vivo* for months [35], [38]. Despite the many advantages of carbon fibers, many labs find manual fabrication of these arrays to be arduous. Some groups [39] combine carbon fibers into bundles that collectively result in a larger (^~^200 μm) diameter than a traditional microwire [39], [40]. Others have fabricated individuated carbon fiber electrode arrays, though their methods require cleanroom-fabricated carbon fiber guides [41]–[43] and equipment to populate their arrays [35], [42], [43]. To address this, we propose a method of fabricating a carbon fiber array that can be performed at the lab benchtop that allows for impromptu modifications. The resulting array maintains individuated electrode tips without specialized fiber populating tools. Additionally, multiple geometries are presented to match the needs of the research experiment. Building from previous work [35], [41], [44], this paper provides detailed methodologies to build and modify several styles of arrays manually with minimal cleanroom training time needed.

## Methods

### Carbon Fiber Array Assembly

Carbon fiber arrays are composed of three parts: a custom printed circuit board (PCB), a backend connector, and an inexpensive sample of 6.8 μm carbon fibers (T-650/35 3K, Cytec Thornel, Woodland Park, NJ). All design files associated with designs presented below are available for download, including three different PCBs: “Flex Array”, “Wide Board”, and “ZIF” (Figure 1) on the MINT website (https://mint.engin.umich.edu/technology-platforms/carbon-fiber-electrodes/). The population and functionalization of a 16-channel Flex Array build is described in detail (build video in supplemental). Several tip optimization techniques to improve electrophysiological recording will also be discussed. A complete materials list including cost is shown in Table 1, with processing steps explained below.

**Figure 1:**
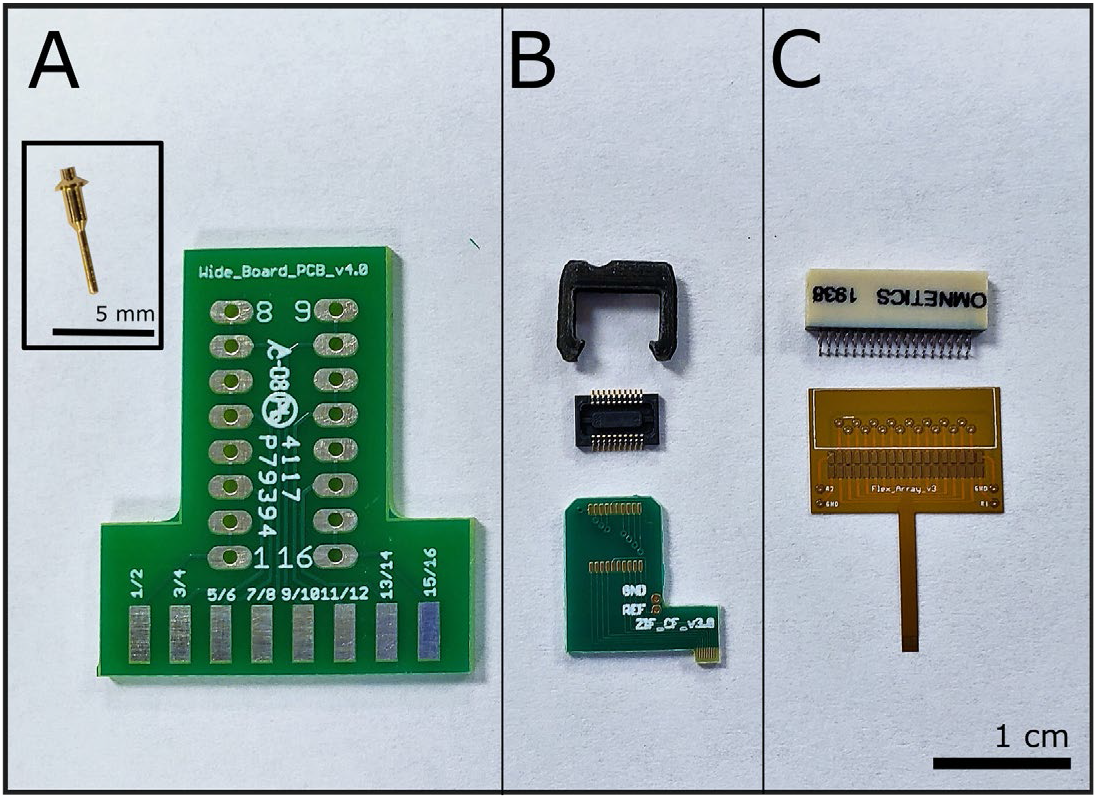
Connectors and associated printed circuit boards. (A) Wide Board with one of sixteen necessary connectors in inset. (B) ZIF and one of two Hirose connectors and TDT shroud. (C) Flex Array with 36-pin Omnetics connector.

**Table 1:**
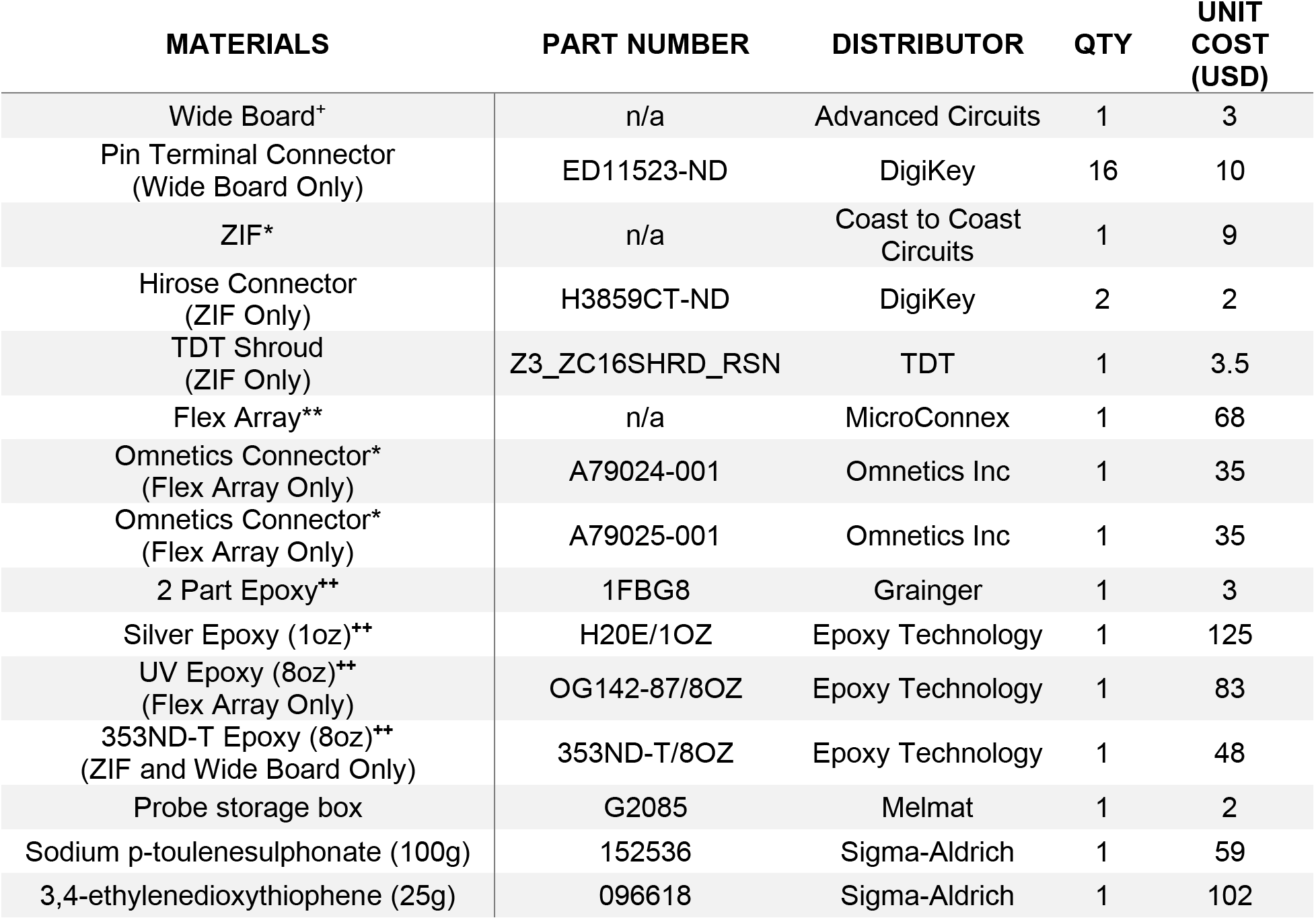
Estimated prices for one board based on 2020 prices. These prices are based on publicly available price listing and do not take into account academic pricing. *Assumes an order of 100, **Assumes an order of 50 with initial $800 NRE charge, +Assumes an order of 200, ++Price is for initial purchase, but can be used for multiple builds.

### Printed Circuit Boards

Wide Board, a ZIF based PCB (referred to as ‘ZIF’ here onwards), and Flex Array PCBs were designed in Eagle CAD (Autodesk, San Rafael, CA). Wide Board and ZIF designs were commercially manufactured (Advanced Circuits, Aurora, CO) and are compatible with Tucker-Davis Technologies (TDT) headstages (Figures 1 A and B respectively). Flex Arrays were fabricated at a commercial facility (MicroConnex, Snoqualmie, WA) (Figure 1 C). Soldering pad and trace sizing vary between each design (Table 2). Wide Boards are the easiest to fabricate. They have a pitch of 3 mm, exposed trace size of 1.5 mm x 4 mm, and are useful for applications where interelectrode distance doesn’t matter, for example, soak testing or testing coating viability. The 16-channel ZIFs have a pitch of 150 μm and an exposed trace size of 0.75 mm x 0.07 mm, which is sufficiently small for insertion testing or acute or chronic ePhys recordings. The smallest of these three designs is the 16-channel Flex Array, with an electrode pitch of 132 μm. Due to the small pitch, two traces are used per fiber to help align the fibers and create a well for the silver epoxy. One fiber per trace is possible (66 μm pitch, for 32-channels) with smaller particle epoxy, but requires a skilled hand to place the epoxy and fiber without shorting the electrodes.

**Table 2:** Each PCB has a different connector and pitch associated with it.

| PCB NAME | CONNECTOR | SOLDERING<br>PAD SIZE<br>(mm) | EXPOSED<br>TRACE SIZE<br>(mm) | TRACE<br>PITCH<br>(μm) | CHANNELS |
| --- | --- | --- | --- | --- | --- |
| Wide Board | Mill-Max<br>9976-0-00-15-00-00-03-0 | 3.25 x 1.6 | 1.5 x 4.0 | 3000 | 8 |
| ZIF | Hirose<br>DF30FC-20DS-0.4V, | 0.23 x 0.7 | 0.75 x 0.07 | 152.4 | 16 |
| Flex Array | Omnetics A79024-001 | 0.4 x 0.8 | 0.6 x 0.033 | 132 | 16 |

### Soldering Omnetics

The first step in building any of these devices is soldering the connector. This requires the use of a stereoscope (SMZ445, Nikon Instruments Inc, Melville, NY) and a soldering iron with a fine tip (0.1-0.2mm). For a lab without soldering equipment, this step can be outsourced to any PCB assembly house. Due to the melting temperature of the polyimide board, a soldering iron temperature of 315°C (600°F) was used to reduce the chance of pads separating from the board. Flux was applied to all contacts before a small amount of solder was placed on the back row of pads. Solder mounds had flat tops so the Omnetics pins were able to sit evenly across them (Figure 2 A). The two outer-most pins were pushed into the solder with the tip of the iron to secure the connector in place. The remaining pins were secured by pushing the tip of the iron between the front pins and pushing them down (Figure 2 B). The front pins were soldered to their respective pads. The remaining flux was cleaned off with 100% isopropyl alcohol (IPA) rinses and a brush (855-5, MG Chemicals, Canada) with bristles cut down to ^~^5 mm.

**Figure 2:**
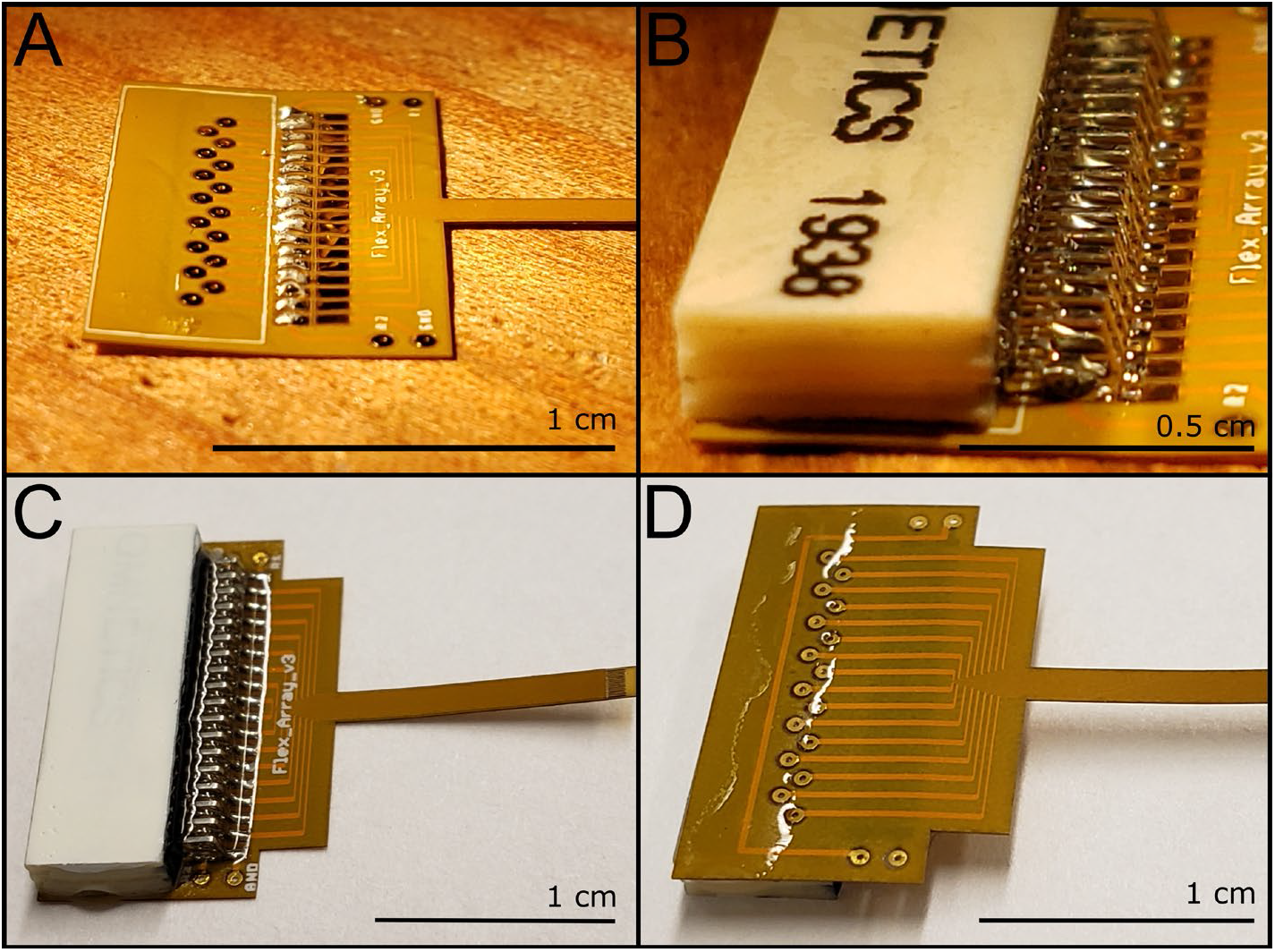
Soldering and insulation steps for the Flex Array. (A) Laying the solder for the bottom Omnetics pins. (B) Back pins secured in place with the front pins ready for soldering. (C) Epoxy insulated Flex Array, note that the epoxy does not cover the reference and ground vias on either side, and (D) Back side of the Flex Array with a band of epoxy across the pad vias (not the ground and reference vias) and wrapped around the side of the board toward the edge of the Omnetics connector.

To prevent the Omnetics connector from deforming and pulling away from the Flex Array, the connections were insulated using a two-part epoxy (Sy-SS, Super Glue Corporation, Ontario, CA). Epoxy was mixed in a dish, pulled into a 1 mL syringe, excess epoxy was wiped from the tip, and a 23 G needle attached. Epoxy was applied bevel side down against the top of the pins to encase the pins and minimizing air bubbles (Figure 2 C). A small amount of epoxy was applied to each side of the Omnetics connector on the board to secure the two during future handling steps (Figure 2 D). Boards were left to cure overnight at room temperature.

### Fiber Placement and UV epoxy

The prepared PCB was placed onto putty under the stereoscope (in the video, the putty is placed on a wooden block for ease of movement). Pulled glass capillaries (TW120-3, World Precision Instruments, Sarasota, FL) were made using a glass puller and filament (P-97 and FB315B, Sutter Instrument Co., Novato, CA) under the following settings: Heat= 900, Pull= 70, Velocity= 35, Time= 200, Pressure= 900 (numbers are unitless and specific to this device). Pulled capillary tips were cut to easily fit between the traces of the board (Figure 3 A). Silver epoxy (H20E, Epoxy Technology, Billerica, MA) was mixed in a dish according to manufacturer specifications. The glass capillary tip was dipped into silver epoxy and applied between pairs of adjacent traces (Figure 3 B) resulting in 8 pairs of connected traces. Traces are shorted together in this way to ease the manual demand of epoxy and fiber laying, however, one fiber per trace is possible for a practiced user (supplemental).

**Figure 3:**
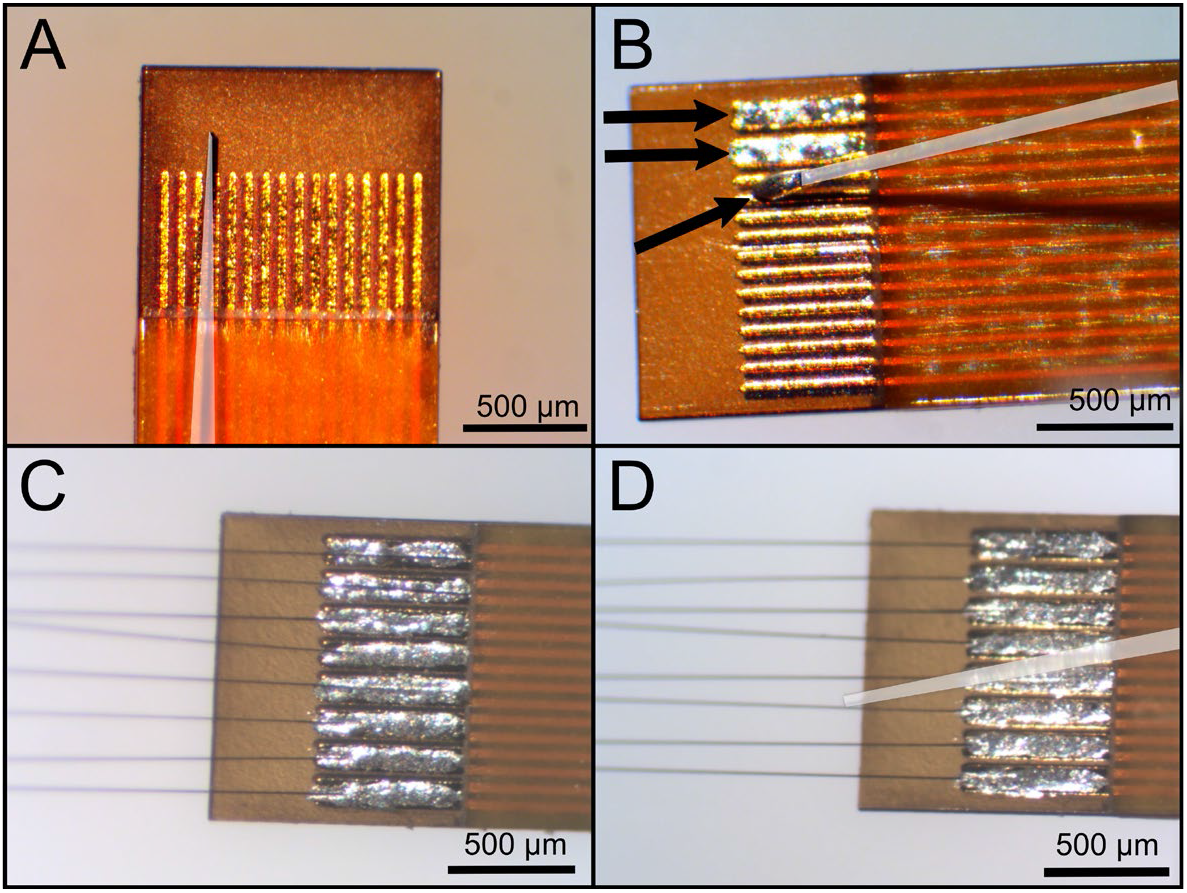
Applying silver epoxy and aligning carbon fibers in between the traces of the Flex Array. Capillaries have been highlighted with a white overlay. (A) The end of the capillary fits between the traces in order to get (B) clean silver epoxy (denoted with arrows at end of the capillary and within traces) deposition without spillover outside of trace pairs. (C) Carbon fibers are placed into the epoxy and then (D) straightened with a clean capillary.

Fibers were initially cut to 2-4 mm in length with a straight razor and separated into single fibers. This was accomplished by gently pulling a laminated piece of paper over the top of the carbon fiber bundle. The laminated paper helps to transfer static into the fibers causing them to separate on their own. A pair of Teflon coated tweezers (11626-11, Fine Science Tools, Foster City, CA) was used to pick up a single carbon fiber segment. Fibers were placed in the silver epoxy mounds (Figure 3 C). A clean capillary was used to adjust the fibers, so they were perpendicular to the end of the board, parallel to the length of the board, and buried beneath the epoxy (Figure 3 D). Carbon fibers were kept clean of silver epoxy past the edge of the board. Arrays were placed on a wooden block without putty, with the carbon fiber portion stick over the edge, and then put into an oven for 20 minutes at 140 °C to cure the epoxy. The wooden block allows for easy transport of the device in and out of the oven, while also holding no static charge that could deform the carbon fibers’ placement. The technique was repeated on the backside of the array resulting in a 16-fiber array. After curing, traces were visually inspected to ensure the connections had no shorts between fibers. Any epoxy shorts or spills were removed with a clean pulled glass capillary. A practiced user can achieve placement angles that are within 0.35 degrees for all fibers perpendicular to the edge of the board [41].

Next, the traces were insulated with a small amount of UV epoxy (OG-143, Epoxy Technology, Billerica, MA) placed on the end of the board using a clean pulled glass capillary (Figure 4 A). The amount of UV epoxy was enough to cover all traces and encapsulate all silver epoxy as this epoxy is meant to insulate the traces and fibers both from each other and from fluid interferences introduced in experiments. The probe was cured under a UV light (SpotCure-B6, Kinetic Instruments Inc, Bethel, CT) for a minimum of 2 minutes (Figure 4 B). The epoxy was checked by lightly tapping the surface with a clean pulled glass capillary to make sure it was fully cured (hard) before repeating on the other side; if not fully cured (sticky, soft), it was placed under the UV light for an additional 2 minutes. Once cured, fibers were cut to 1 mm lengths using stainless steel microsurgical scissors (15002-08, Fine Science Tools, Foster City, CA). When properly insulated, the board will have a small hard bubble on either side (Figure 4 B inset).

**Figure 4:**
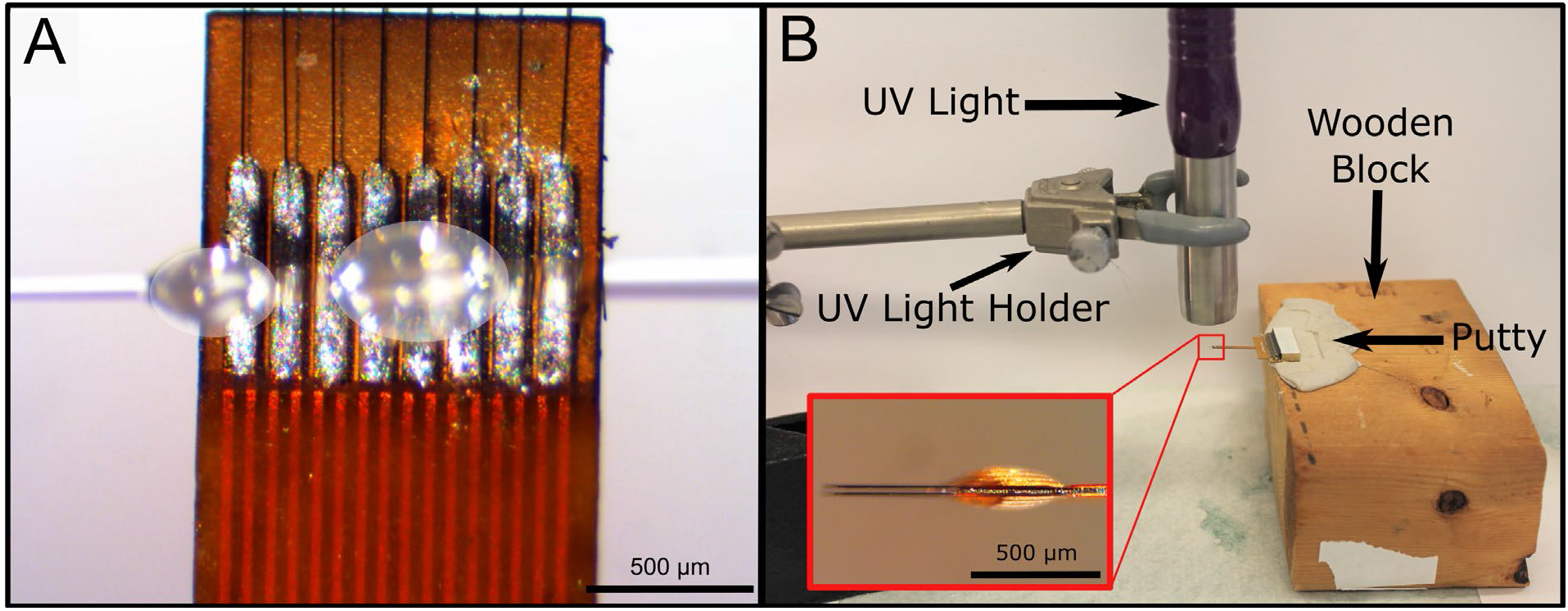
(A) UV epoxy is applied using a clean capillary and two drops of UV epoxy (marked with white overlays). UV epoxy is applied in droplets of 0.25-0.75 mm diameters until the UV epoxy forms a smooth bubble over the top of the traces. (B) Epoxy is cured under a UV light. The Flex Array is placed in putty on a wooden block for ease of movement and alignment underneath the UV light. The UV light is held with a holder about 1 cm above the end of the Flex Array. Inset (B) shows the side profile of a properly insulated Flex Array. The bubble on either side of the board is roughly 50 μm in height.

### Checking Electrical Connections

A 1 kHz impedance scan was taken to confirm the fibers were electrically connected to the Omnetics connector and no shorts existed between fibers. Fibers were submerged 1 mm in 1x PBS (BP3994, Fisher, Waltham, MA) with an Ag|AgCl reference electrode (RE-5B, BASi, West Lafayette, MA) and a stainless-steel rod as the counter electrode. A PGSTAT12 Autolab (EcoChemie, Utrecht, Netherlands) and NOVA software provided by the vendor were used to take the measurements. Results were analyzed using custom MATLAB scripts (MathWorks, Natick, MA). Measurements were taken at multiple steps during the build process to verify connections. Typical impedance ranges varied depending on the build step (Table 3).

**Table 3:** Typical range of impedances after each build stage (n=272) *n=16. PEDOT:pTS treated probes above 110kΩ may still record signals, however all treated electrodes typically fall under this value.

| BUILD STEP | EXPECTED 1KHZ IMPEDANCE (kΩ) |
| --- | --- |
| Bare Fiber | 150-300 |
| Bare Fiber with UV Insulation | 400-500 |
| Parylene-C Insulated Fibers | >50,000 |
| Nd:YAG Laser Cut | <15,000 |
| Blowtorched | 300-400 |
| UV Laser Cut* | 300-500 |
| PEDOT:pTS Coated | <110 |

Once there was confidence in each build step, the number of impedance scans were reduced. Currently, they are performed only prior to parylene-c insulation and then as prescribed by the tip treatment procedure.

### Parylene-C Insulation

The Flex Array’s backend connector was masked using the mating connector (A79025-001) to prevent the internal pins of the Omnetics connector from being coated during the insulation process. A batch of arrays (8-12) were placed into a box with a raised, adhesive platform such that the connector end of the probe was resting on the platform and the majority of the board was overhanging the edge of the raised platform (Figure 5). We used inverted Fisher Tape super glued to a piece of foam as the raised platform.

**Figure 5:**
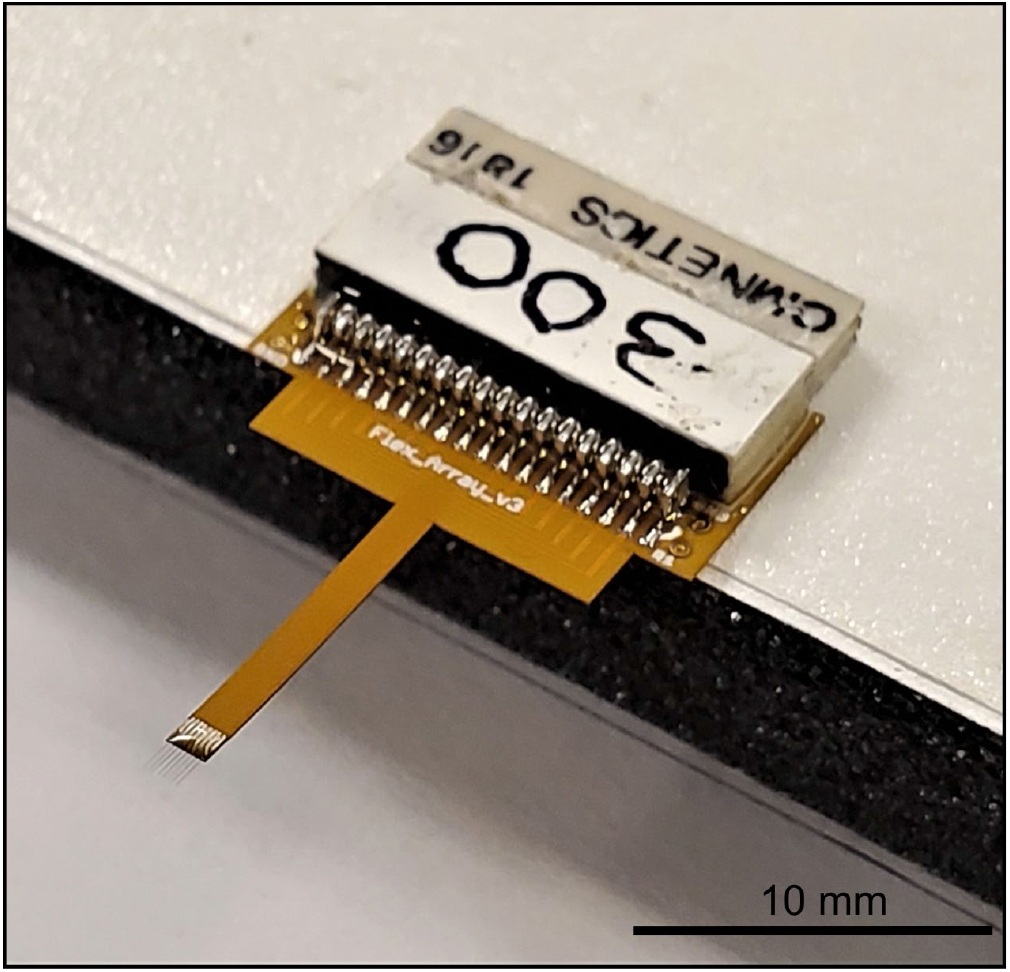
Flex Array prepared for parylene-c coating. The Flex Array is secured during the coating process to a raised foam platform with tape, adhesive side up.

Arrays were coated with a conformal layer of parylene-c (thickness = 800 nm) using the Parylene-c Deposition System 2035 (Specialty Coatings Systems, Indianapolis, IN) located within the Lurie Nanofabrication Facility at the University of Michigan, following deposition rate guidelines of the machine. Many cleanrooms at research universities will have this deposition capability, which is easy for an individual to learn. A batch of 5 probes were sent to Specialty Coating Systems (Indianapolis, IN) to determine the viability of outsourcing this step to remove the only fabrication step that requires a cleanroom.

After parylene-c insulation, the backend masking was removed and the arrays were placed into a new box. A new box is required as the tape in the box that went through parylene-c deposition will be coated as well and unable to hold the arrays in place. The arrays were stored in a cool, dry, and dark place and considered shelf stable. An inventory of arrays was built up and used when needed for experiments.

### Tip Preparation Methods

One of three methods was used to re-expose carbon at the tip of the fiber: Nd:YAG laser cut, blowtorch, or ultraviolet (UV) laser. The fibers must be cut by one of these three methods as scissor cutting alone is not sufficient to reliably re-expose the tip of the carbon fiber [44]. Fibers under 500 μm “self-insert” (require no additional or specialized insertion techniques) into the cortex [41], but for nerve or muscle a final length of ≤ 300 μm with sharpened tips was required [45].

#### Nd:YAG Laser Cut

Fibers were first cut to 550 μm with surgical scissors. A 532 nm Nd:YAG pulsed laser (LCS-1, New Wave Research, Fremont, CA: 5 mJ/pulse, 5ns duration, 900 mW) was used to further expose the carbon in conjunction with a Karl Suss probe station (LC3, SUSS MicroTec, Garching, Germany) for fiber alignment as shown previously [44]. The fibers were aligned inside of a 22 x 50 μm cutting window and the tips were cut off with 2-3 pulses resulting in a final length of 500 μm. The parylene-c ablated only slightly back (<10 μm) from the tip after each cut [44].

#### Blowtorch

While Nd:YAG laser cut fibers reliably re-expose fiber tips, access to such a laser can be limiting. It also only provides blunt cylindrical electrode tips that have some difficulty inserting into muscle and nerve. Thus, a modified approach to previous sharpening methodology [40], [46] was taken using a butane blowtorch (Microtorch MT-51B, Master Appliance, Racine, WI). Fibers were cut to 300 μm using surgical scissors. Using previously developed methods for nerve electrodes [47] an array was submerged in a dish of deionized water with the connector secured to the base of the dish with putty. The board was visually leveled and the water level was adjusted using a pipette and a pen camera (MS100, Teslong, Shenzhen, China) to ensure that the fibers were touching the surface of the water. A 3-5 mm flame from the blowtorch was run over the top of the fibers in a back-and-forth motion to sharpen the fibers (supplemental for video). The array was removed from the putty and inspected under a stereoscope for pointed tips. The process was repeated until points were able to be observed under a stereoscope.

#### UV Laser Cut

A UV laser can also be used to both cut and sharpen carbon fibers similarly to the blowtorch method. While the UV laser is currently unable to be used with Flex Arrays due to the board’s small pitch size between fibers and rows of fibers, it does show promise with the larger pitch of the ZIF and Wide Board designs. This method is being developed to give a pathway to laser cutting with an easily obtainable UV laser to remove the access barrier that the Nd:YAG laser may provide. Thus, carbon fibers (2 mm length) were mounted on a ZIF and parylene-c insulated. A 1500 mW UV laser head (WER, Shanghai City, China) was affixed to three orthogonally configured motorized stages and then its focal point was moved across the fiber plane to cut the fibers to 500 μm [48].

### PEDOT:pTS Coating

For all tip cutting methods, an additional conductive layer must be added to the exposed carbon site to further reduce its impedance. In previous work, poly(3,4-ethylenedioxythiophene):sodium p-toulenesulfonate (PEDOT:pTS) has been used. A 50 mL solution of 0.01 M 3,4-ethylenedioxythiophene (483028, Sigma-Aldrich, St. Louis, MO) and 0.1 M sodium p-toluenesulfonate (152536, Sigma-Aldrich, St. Louis, MO) was stirred overnight, then refrigerated, and replaced every 30 days. This solution was stored in a light resistant container as it is light sensitive.

Probe impedances were taken in 1x PBS solution with the same parameters used previously; “broken” (missing fiber) and “bad” (impedances > 1MΩ) channels were noted and not included in the PEDOT:pTS coating. Fibers with a good connection (typically 14-16 of the fibers) were electroplated with the PEDOT:pTS solution by applying 600 pA per fiber for 600 s using a PGSTAT12 Autolab. After electroplating, a final impedance measurement was taken and fibers with an impedance over 110 kΩ were designated “bad” in the probe’s documentation.

### Finalizing the Probe

The final step for finishing the probe was to solder ground and reference wires (Teflon Coated Silver Wire #AGT05100, World Precision Instruments, Sarasota, FL) to the probes. As the ground and reference vias were coated in parylene-c, they were scraped clean with tweezers on the top and bottom of the board. Two 5 cm silver wires were de-insulated on either end (1 cm and ^~^2 mm). The 2 mm exposed portion of the wires were placed into the ground and reference vias and soldered into place. The excess wire was cut away from the backside solder joint (Figures 6 A and B). Probes were placed in a storage box with the reference and ground wires secured away from the electrode tips (Figure 6 C).

**Figure 6:**
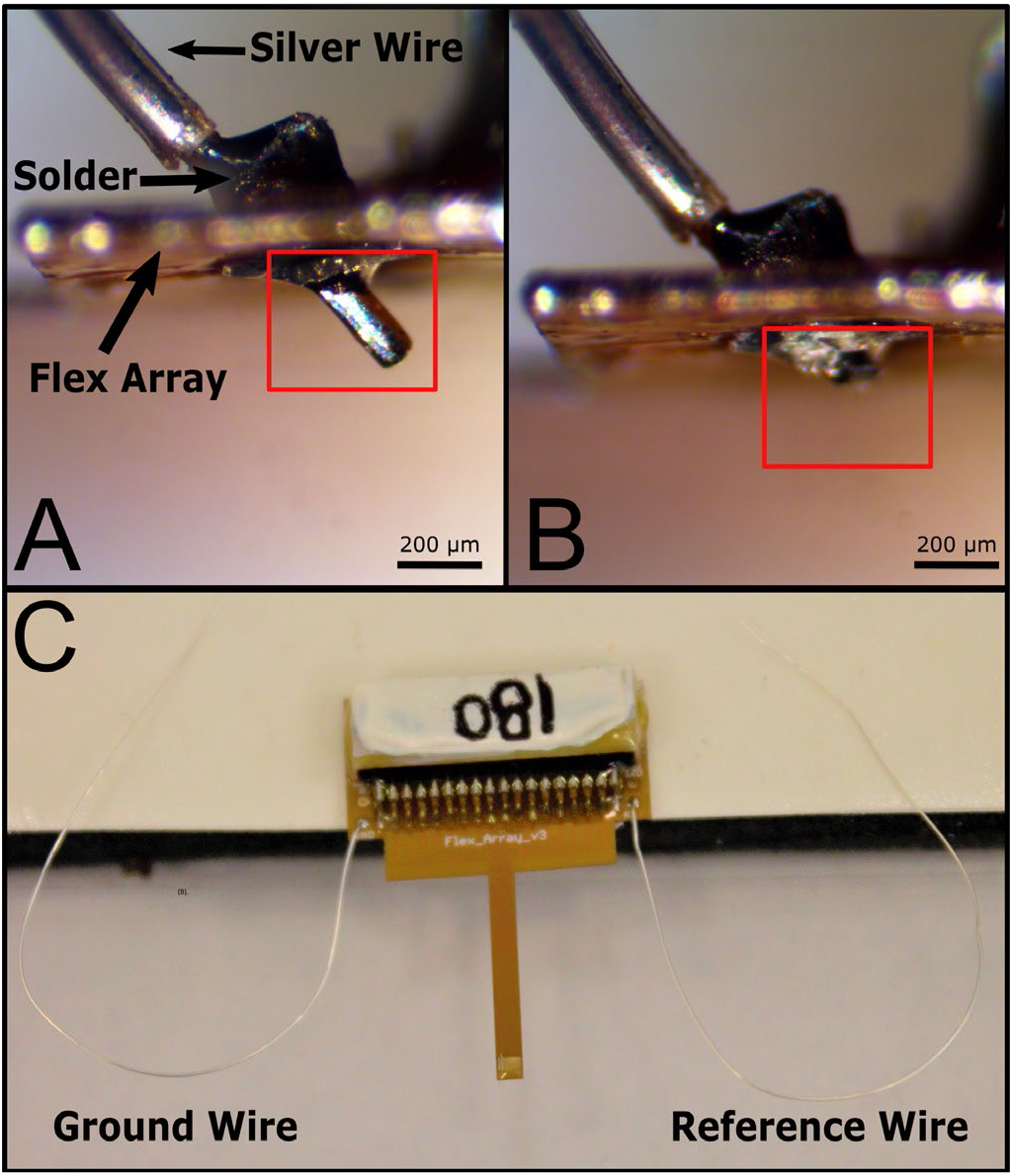
Ground and reference wires attached to the finalized Flex Array. Solder applied to each side of the via (A) and excess wire (red boxes) is (B) removed from the backside of the probe. Final Flex Array stored for future use.

### Surgery Protocol

All animal procedures were approved by the University of Michigan Institutional Animal Care and Use Committee.

Surgical procedures for acute recordings followed Patel et al 2015 [41]. To summarize, an adult male Long Evans rat weighing approximately 300 g was anesthetized using a combination of ketamine (90 mg/kg) and xylazine (10 mg/kg). A bone screw (19010-00, Fine Science Tools, Foster City, CA) was used as the common reference and ground at the posterior edge of the skull. A 2.5 mm by 2.5 mm craniotomy was made over the right hemisphere’s motor cortex. After dura resection, a ZIF array with 4 UV laser treated fibers was inserted to a depth of 1.2 mm. All ePhys data was collected using a ZC16 headstage, RA16PA pre-amplifier, and RX5 Pentusa base station (Tucker-Davis Technologies, Alachua, FL). The pre-amplifier high-pass filtered at 2.2 Hz, anti-alias filtered at 7.5 kHz, and sampled at 25 kHz. The recording session was 10 minutes long.

### Spike Sorting

Offline Sorter (OFS, Plexon, Dallas, TX) software was used to spike sort the data following the methods outlined in [49]. Channels were high-pass filtered (250 Hz corner, 4^th^ order Butterworth) and waveforms were detected at −3.5*RMS threshold. Cluster centers were identified in principle component states using a K-means sorting method. Obvious noise clusters were eliminated from the data set. A Gaussian model was used to cluster the remaining clusters. Spikes with similar characteristics were combined and averaged over the cluster. Carbon fiber electrodes with discernible units were deemed viable. A minimum of 10 waveforms were required for a unit to be included in the data.

### SEM Imaging

An FEI Nova 200 Nanolab Focused Ion Beam Workstation and Scanning Electron Microscope (FEI, Hillsboro, OR) was used for SEM imaging of Nd:YAG laser and blowtorch prepared fibers. Prior to imaging, samples were gold sputter coated with an SPI-Module Sputter Coater (SPI Supplies, West Chester, PA). Images of UV laser prepared fibers were obtained with the JEOL InTouchScope Scanning Electrode Microscope (JSM-IT500HR, JEOL, Tokyo, Japan).

## Results

### Tip Validation: SEM Images

Previous work [44] showed that scissor cutting resulted in unreliable impedances as parylene-c folded across the recording site. Scissor cutting is used here only to cut fibers to a desired length before processing with an additional finish cutting method. SEM images of the tips were used to determine exposed carbon length and tip geometry (Figure 7).

**Figure 7:**
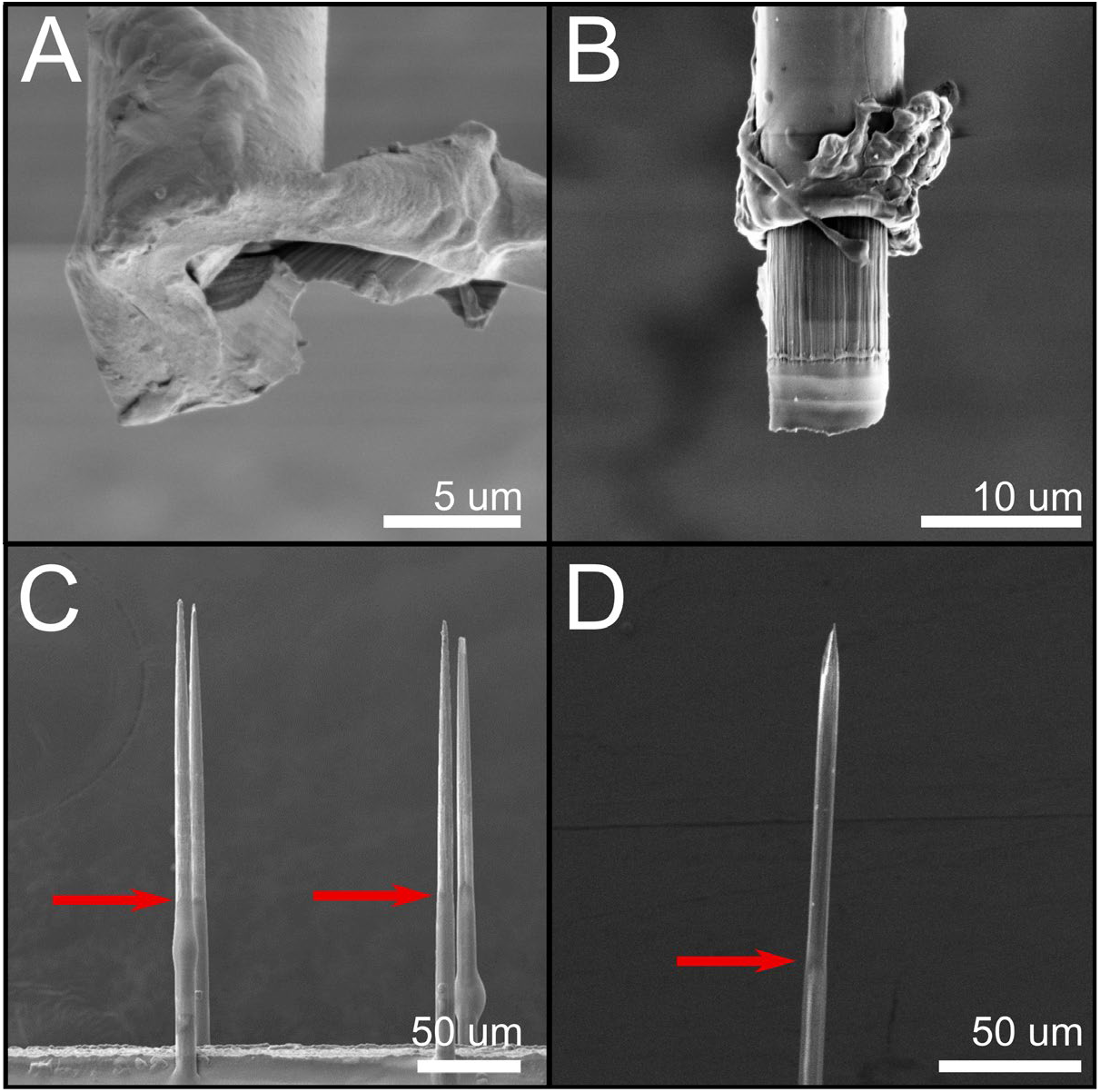
SEM images of fibers with different tip cutting techniques. (A) Scissor cut fiber with very little exposed carbon. (B) Nd:YAG laser cut. (C) Blowtorched fiber with ^~^140 μm of carbon exposed from the tip. (D) UV laser cut fibers with ^~^120 μm of carbon exposed from the tip. Red arrows indicate the transition area between parylene and bare carbon fiber.

Scissor and Nd:YAG laser cut fibers were previously reviewed [35], [44]. Scissor cut fibers (Figure 7 A) have inconsistent tip geometries with parylene-c folding over the end when cut [44]. The Nd:YAG laser cut fibers remain consistent in recording site area, shape, and impedance (Figure 7 B). Blowtorched fibers [47] lead to the highest electrode size and shape variability, but also resulted in a sharpened tip allowing for insertion into tough tissue and on average 140 μm of carbon was re-exposed with a smooth transition area between the carbon and parylene-c insulation. UV laser cut fibers were similar to with blowtorched fibers, showing 120 μm of carbon exposed from the tip. Impedances indicated that either the UV laser or blowtorch tip cutting methods are suitable for ePhys and are viable solutions for labs without access to an Nd:YAG laser.

### Tip Validation: Electrical Recording

Figure 8 shows resulting impedances from each preparation method using Flex Arrays. The resultant values are within an appropriate range for ePhys recording. Nd:YAG laser cut fibers resulted in the smallest surface area but the highest impedances, even with the PEDOT:pTS coating (bare carbon: 4138 ± 1438 kΩ; with PEDOT:pTS: 27 ± 18 kΩ; n = 262). This is followed by the inverse relationship in blowtorched (bare carbon: 308 ± 121 kΩ; with PEDOT:pTS: 16 ± 13 kΩ; n = 262) and UV Laser cut (bare carbon: 468 ± 342 kΩ; with PEDOT:pTS: 27 ± 8 kΩ; n = 7) fibers that have a large surface area and low impedances. However, in all cases, the PEDOT:pTS coated fibers do fall under the 110 kΩ threshold that was set previously to indicate a good, low impedance electrode.

**Figure 8:**
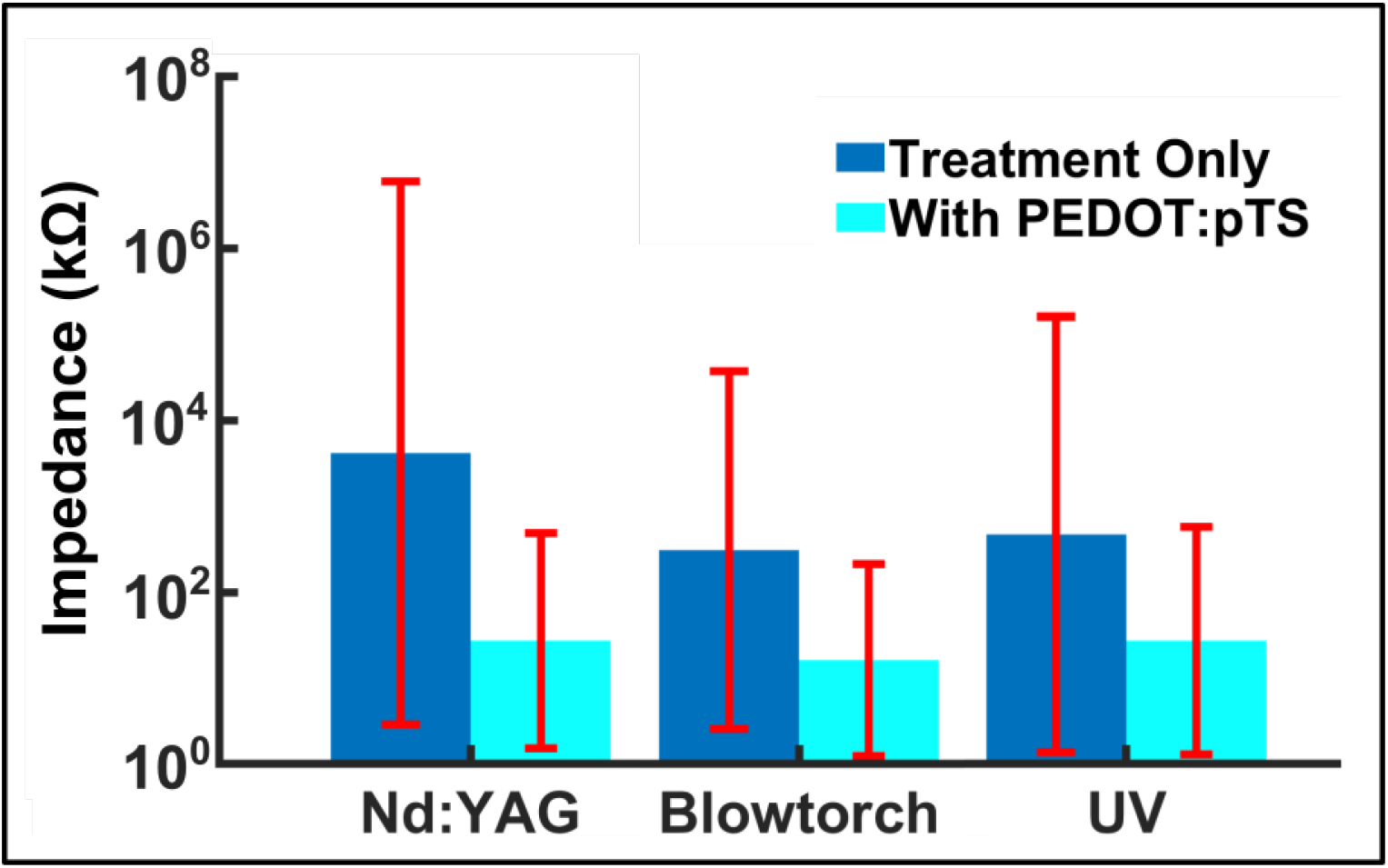
Impedance (mean ± standard deviation) differences between only applying the treatment (bare carbon exposed) and with the addition of PEDOT:pTS. In all cases the addition of PEDOT:pTS decreases the impedance by an order of magnitude.

Acute ePhys recordings were taken from a Long Evans rat acutely implanted with a ZIF array with UV laser cut and PEDOT:pTS treated fibers to demonstrate the viability of this method. EPhys has previously been tested and proven with scissor cut [44], Nd:YAG [35], and blowtorch treated fibers [45], [49]. Acute recordings from four UV laser treatment fibers that were simultaneously implanted in rat motor cortex (n=1) are presented in Figure 9. Three units were found across all fibers suggesting that the treatment of the fibers with the inexpensive UV laser is similar to other cutting methods that enable the carbon fiber to record neural units, as would be expected by the SEMs and impedances.

**Figure 9:**
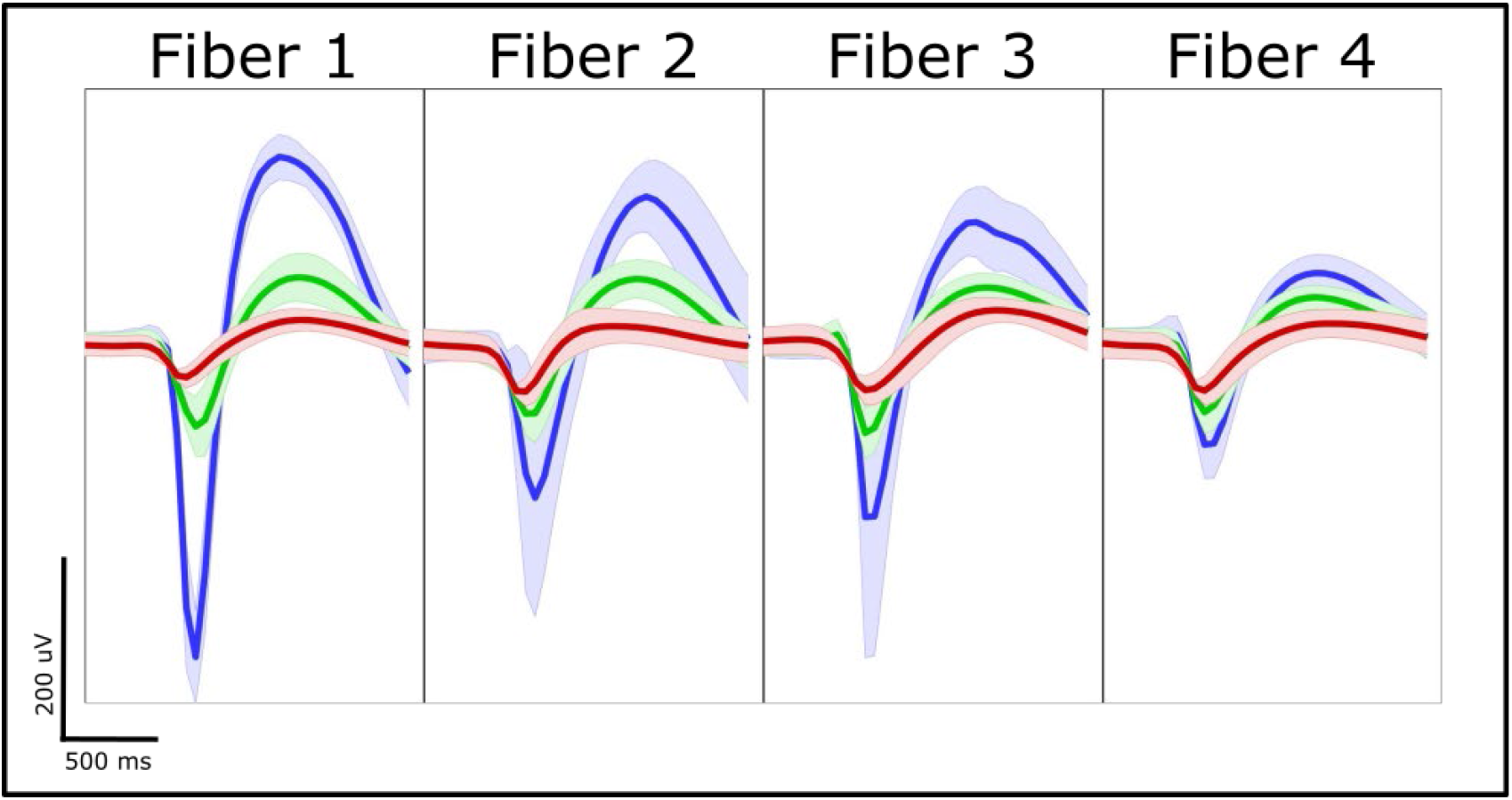
Acute electrophysiological spiking data from four UV laser cut electrodes.

While carbon fiber arrays are easily built and modified to suit the user’s needs, it should be noted that additional validation is necessary for some builds (Table 4) while others are less suitable for certain end tasks.

**Table 4:** Validated uses of each board with the cutting methods described. All cutting methods included electrodeposition of PEDOT:pTS. ‘Not Viable’ indicates that a form factor of the design prevents this tip treatment from being tested at this time (i.e. fiber pitch).

| PREPARATION METHOD | WIDE BOARD | ZIF | FLEX ARRAY |
| --- | --- | --- | --- |
| **Nd:YAG** | Impedance, SEM, acute ePhys | Impedance, SEM, acute/chronic ePhys | Impedance, SEM, acute/chronic ePhys |
| **Blowtorch** | Impedance, SEM, acute ePhys | Impedance, SEM, acute/chronic ePhys | Impedance, SEM, acute/chronic ePhys |
| **UV Laser** | Not yet validated | Impedance, SEM, acute/chronic ePhys | Not Viable |

### Commercial Parylene-c

Commercially coated arrays were determined to have a parylene-c thickness of 710 nm by the vendor, well within the target range of insulation. The arrays were prepared for ePhys recordings using the blowtorch tip preparation. Impedances were taken after preparation of the tips and compared to existing data. A blowtorched and PEDOT:pTS coated probe had an average of 14.5 ± 1.3 kΩ impedance across 16 fibers. SEM images were taken of the tip and shank to compare parylene-c deposition (Figures 10 A and B, respectively). These results show the use of a commercial vendor did not change expected impedance values, suggesting that this will be an equally viable substitution to deposition at the university cleanroom.

**Figure 10:**
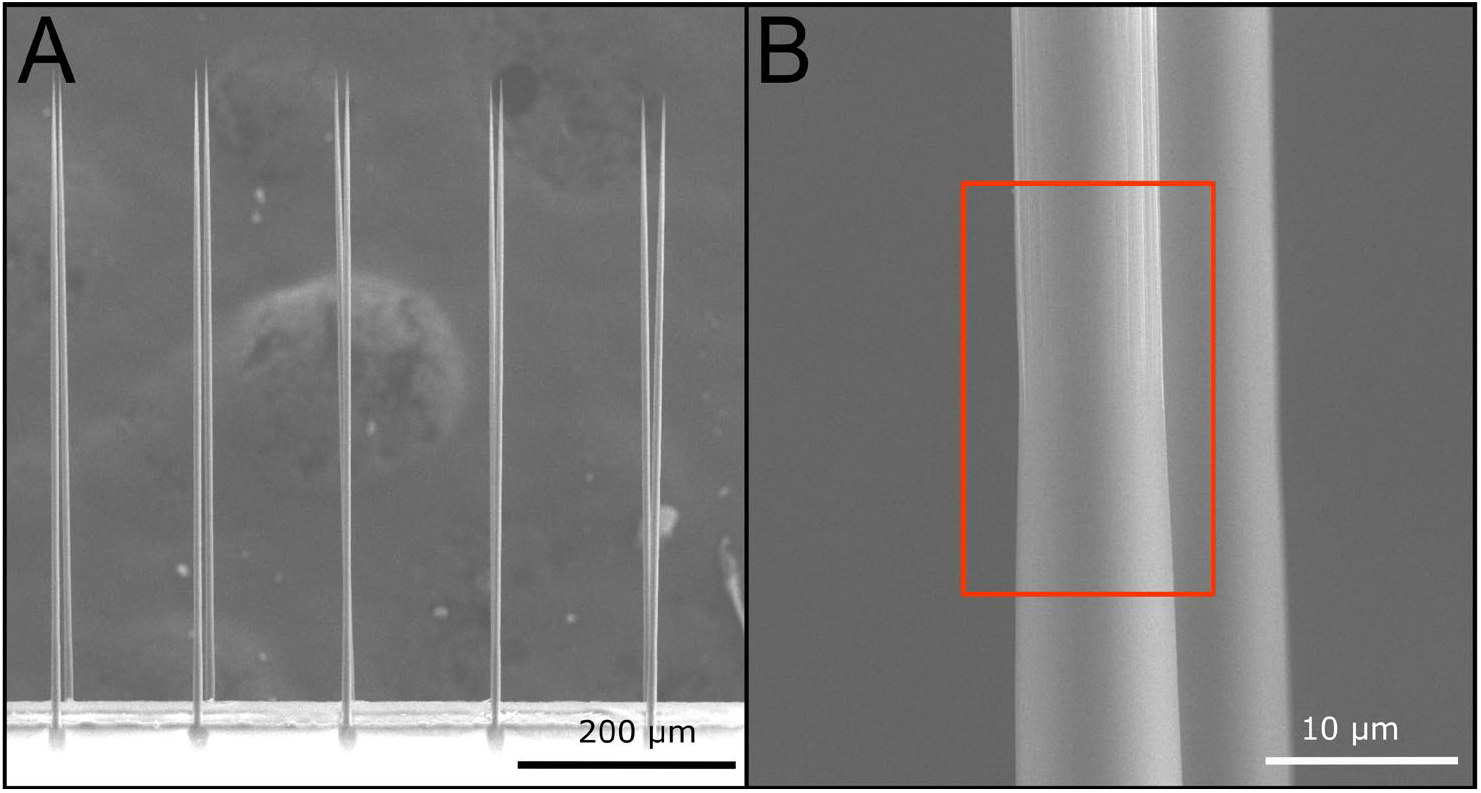
Commercial parylene-c coated arrays. (A) The sharpened array shows uniform sharpening across all fibers indicating that there are no drawbacks to commercial coating. (B) The transition (red box) between bare carbon fiber and parylene-c after blowtorching shows no discernable difference between arrays coated in a cleanroom facility.

### Device Cost Analysis

Provided all tools and bulk materials (epoxies, solder, etc.) are accessible to the researcher, a parylene-c user fee of $41, and a batch of 8 probes, the total materials cost is $1168 ($146 per probe). Personnel effort (Table 5) is around 25 hours for the batch. If using a substituted fabrication step, the cost of the probes will vary based on commercial parylene-c coating cost ($500-800 quoted). The time for build steps (Table 5) is grouped together for all instances of a repeated tasks for simplicity. Build times for designs with a larger pitch (Wide Board and ZIF) are dramatically reduced as the manually intensive steps (e.g. carbon fiber placement) are easier and faster to complete.

**Table 5:**
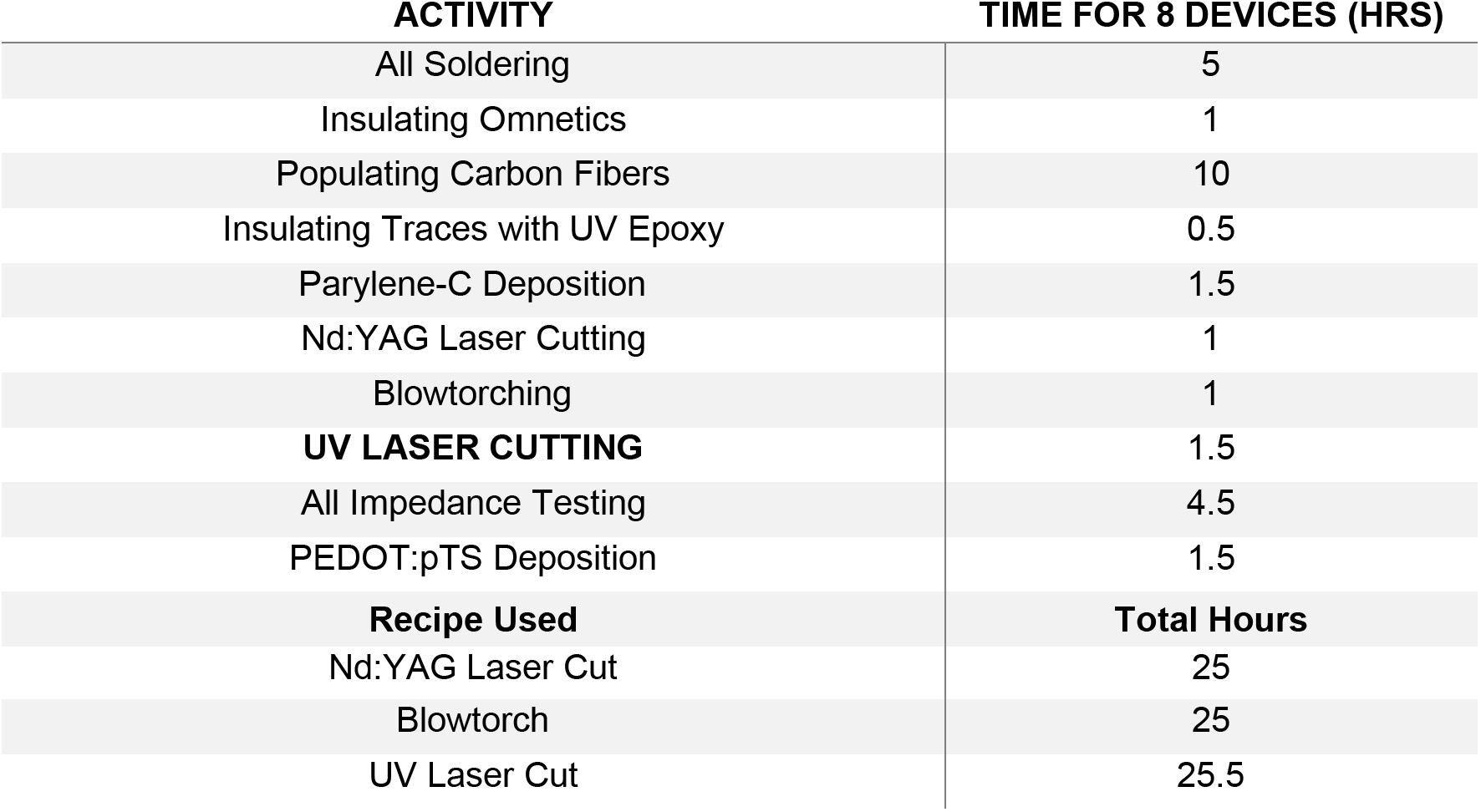
Time required for each step of a fabrication process. Soldering of the Omnetics and ground and reference wires have been combined here to simplify the activity list.

## Discussion

### Troubleshooting Build Issues

Silver epoxy deposition tends to fail for several reasons: the width of the capillary is too wide to fit between traces, the width of the capillary is too thin to pick up and deposit epoxy, or an excess of epoxy is on the capillary. The first two problems can be solved by cutting a new capillary that is a more appropriate size; the latter by dipping the capillary into the epoxy with a lighter hand or removing a portion of the epoxy blob by gently dabbing the capillary onto a spare nitrile glove.

Deciding how to prepare the electrode is often a difficult decision for many users. However, determining what is needed for the experiment will help illuminate the decision. For acute surgeries, blunt tips can be used if the site size of the electrode is important, however they will only insert into softer tissue (brain) and only at sub-500 μm target depths. Going into deeper brain structures is possible using a glass cannula[50], but this can cause scarring and associated unreliability in ePhys recordings. Fibers must be less than 300 μm when sharpened to be able to penetrate the nerve tissue as the shorter length provides a stiffer backbone for insertion. Sharpened fibers have also recently been observed to penetrate to 1mm depths in the brain [49].

While the arrays discussed in this paper are an excellent starting point for many labs, newer probes using carbon fibers have also been developed to chronically target deeper areas. The Chestek lab high density carbon fiber array provide users the ability to insert 16 carbon fibers to depths up to 9 mm with minimal scaring [50]. The Maharbiz lab has also developed a higher channel probe that was inserted to depths of 2.5-3 mm [51]. The Cox lab has also developed a carbon fiber bundle probe ^~^1 mm in length [39]. These probe geometries will be of greater use to labs doing chronic work in deeper brain structures.

### Parylene-C Accessibility

Parylene-C is a method of conformal coating at room temperature that has been used as a biocompatible insulator in many implanted devices. The technique requires a specialized tool in a cleanroom and takes about an hour to learn. A cursory survey of institutions that have previously requested carbon fiber arrays from our group was conducted to determine parylene-c deposition accessibility. We found that out of 17 institutes, 41% had access to parylene-c coating systems on their campus. For universities without access to a parylene-c coating system, commercial coating services are a viable alternative as we have demonstrated. Alternatively, outsourcing to a nearby university cleanroom may also be of interest to labs with no direct access to a parylene-c deposition system. To reduce the cost per device, we advise sending out larger batches of arrays as commercial systems are often able to accommodate larger samples.

### Optimizing Tip Preparations

Additional tip preparations need to be investigated for these fibers as the current tip preparations do require the end user to choose between penetrating ability and a small recording site. While the Nd:YAG laser cut fibers provide a small site size [44], the ability to penetrate stiffer tissue (muscle, nerve) is almost non-existent and access to a laser setup capable of this cutting technique can be difficult and expensive. While blowtorching allows for a quick and economical way to get sharpened tips that can penetrate many tissues [45], the tip geometry is large and may be inconsistent from fiber to fiber [47]. UV laser cutting falls to the same issues with the added obstacle of having to engineer a way to align the laser with fibers and current restrictions on using this method with the Flex Array due to the pitch of the fibers being smaller than the laser’s focal point diameter. Previous work showed fabrication of small, sharpened fibers via etching [52], [53]. This approach could result in a small, reliable electrode geometry and would preserve the sharpened tip necessary for penetrating nerve and muscle.

Our current tip coating, PEDOT:pTS, may also need to be replaced as it tends to degrade overtime, which is an undesirable trait for a chronic probe [35], [44], [54]. A lack of PEDOT:pTS longevity, leads to higher impedances and therefore lower signal quality in part due to increased background noise. To increase longevity in our fiber tips, investigation into the feasibility of platinum-iridium coatings is being conducted. Platinum-iridium would allow for a greater surface area [44], [55] concentrated on the tip of the electrode, keeping a low impedance and [55]–[57] allow for longer, chronic stability [55], [57].

Other coatings such as PEDOT/graphene oxide [58] and gold [59] have been utilized to lower carbon fiber electrode impedances, though these coatings are typically used for chemical sensing probes rather than for ePhys recordings. Due to the inherent properties of carbon fibers [60], the carbon fiber array presented here can be converted from a probe optimized for ePhys to a chemical sensing device with a simple change of tip preparation [50].

### Further Experimental Options

The carbon fibers arrays in this paper have also been successfully fabricated using the methods presented here and utilized to create dopamine sensing arrays. This is accomplished by ablating the parylene-c away from the tip of the probe using the Nd:YAG laser, at a lower power setting, and exposing 50 μm of fiber [50]. Using custom hardware and software, simultaneous high density channel recording has recently been achieved [50]. The probe’s ability to be customized for ePhys or dopaminergic recordings offers yet another layer of adaptability to our carbon fiber array. Coupled with the new coating techniques previously mentioned, this opens up a variety of possible probe variants that could be made and modified, quickly and inexpensively.

### Automated Carbon Fiber Placement

The manual population of the carbon fibers is labor intensive and has a steep learning curve. Many new users struggle with the fibers breaking during handling, flying away due to static forces, and seeing the fibers when not under the stereoscope. While these issues can be addressed with some modifications to the work area, the process is still intensive and slow. Automated carbon fiber placement of an array has been achieved in a different probe configuration [42] in the Maharbiz lab. This technology allows for carbon fiber to be inserted and aligned into a probe with the use of a camera, motorized stages, custom guides, and a custom computer algorithm. Modifying this set up would allow for new users to avoid the learning curve of handling small (2-4 mm long) fibers and allow them to manually feed a longer fiber (> 1 cm) into a machine that would then place and align the fibers. If this technology were implemented, probes would be able to be made faster and more accurately with less training required. However, using the approaches discussed in this work, we believe that virtually any lab with experience working under a stereoscope should be able to fabricate their own fully functional carbon fiber electrode arrays.

## Supporting information

Flex Array Carbon Fiber Population

## Acknowledgment

This work was financially supported by National Institutes of Health National Institutes of Neurological Disorders and Stroke (UF1NS107659 and UF1NS115817) and the National Science Foundation (1707316). The authors acknowledge financial support from the University of Michigan College of Engineering and technical support from the Michigan Center for Materials Characterization and the Van Vlack Undergraduate Laboratory. The authors thank Dr Khalil Najafi for use of his Nd:YAG laser and the Laurie Nanofabrication Facility for use of their parylene-c deposition machine.

