## Supplemental Methods for "Benchtop Carbon Fiber Microelectrode Array Fabrication Toolkit"

*Build Differences*

Wide Board

*Soldering:* Wide Board connectors use a DIP connector which utilizes pins that slot into vias on the board. A DIP connector was placed on the back side of the board with the pins of the connector protruding through the vias into the connector, holding them in place for soldering. Flux was applied to the pins and soldered in place. Flux was cleaned with IPA and cotton applicators.

*Fiber Population and Insulation:* PCBs were roughened around the traces at the end of the board to allow for better adhesion of the insulating epoxy in a later step. Silver epoxy was applied directly to the traces with the wooden tip of a cotton applicator and a carbon fiber was placed on each trace. Carbon fibers are typically cut to be much longer (1cm) for these devices as they are used primarily in soak testing [35], [44]. This additional length allows more flexibility within the carbon fibers, so additional care is required when handling the fibers. Epoxy was cured using manufacturer recommended parameters. Traces were insulated with 353ND-T epoxy and cured following manufacturer recommended settings,

*Tip Cutting:* Due to the typically longer length of fibers on these boards, implementing an air dam to reduce currents around the fibers is necessary to keep them still for the Nd:YAG laser. Blowtorching is not a viable solution as the fibers tend to float around and bend rather than extend to the surface of the water. Due to the movement of the motorized stage and difficulty of including an air dam, the UV laser cutting method was not tested.

ZIF

*Soldering:* The ZIF PCB utilizes a Hirose connector designed to interface with Tucker-Davis Technologies ZIF headstages. Flux was applied to the soldering pads on one side of the PCB and the connector was secured onto the board by first soldering the opposite corners of the Hirose connector before soldering the rest of the connector's pins. This was then repeated on the other side of the board. Flux was cleaned off the board with IPA. A plastic shroud was then placed around the end of the board and pressed into the board until the ends of the shrouds 'clicked' into place around the connector. At this point, a drop of super glue was applied to one side of the shroud to secure the shroud to the board, being careful not to get any glue on the soldering pads.

*Fiber Population and Insulation:* Silver epoxy was applied to each trace with a pulled capillary to form a small mound along the length of the trace. Carbon fibers were then placed into the epoxy keeping the fibers straight by aligning with the trace. Epoxy was cured at 140$^{o}$C for 20 minutes then deposition was repeated on the opposite side of the board.

Insulating epoxy (353ND-T, Epoxy Technology, Billerica, MA) was prepared according to vendor specifications and applied to the traces of the boards using a glass capillary. Boards were cured at 120$^{o}$C for 20 minutes. This process was repeated on the opposite side of the board.

32 Channel Flex Array

Instead of shorting pairs of traces on the Flex Array together, one carbon fiber can be attached to one individual trace to provide 32 channels of individual recording data. This method takes months of practice to build up the dexterity needed for this precise technique. Testing on implants and recording with these devices have been limited in our lab due to unforeseen lab closures.

*Fiber Population:* Carbon fibers were picked up with tweezers and the tips of the fiber were dipped into a different silver epoxy (HPS-030LV, NovaCentrix, Austin, TX). The epoxy was distributed along the trace until enough was deposited for the carbon fiber to be placed on the trace and buried in the epoxy. This was repeated for the remaining traces. Fibers were aligned with a clean pulled glass capillary and then cured at 140$^{o}$C for 20 minutes. Fibers were placed on the backside of the board similarly and then insulated with UV epoxy following the standard procedure.
